# RR3DD 2.0: an expanded database for RNA three-dimensional structure classification

**DOI:** 10.64898/2026.11.18.767751

**Authors:** Danyang Li, Xudong Liu, Chengeng Liu, Yuchen Zhu, Shiyong Liu

**Affiliations:** School of Physics, Huazhong University of Science and Technology, Wuhan, Hubei, 430074, China

## Abstract

RNA performs diverse biological functions by folding into specific 3D structures, which are often more conserved than their sequences. Systematic classification of RNA 3D structures is fundamental to understanding structure-function relationships. We previously established RR3DD, the first classification database based on global 3D structural similarity. However, its version 1.0 contained data only up to July 2021 and can no longer reflect the rapidly growing RNA structure space since then. Here, we present RR3DD 2.0. Starting from the 7,466 PDB-format chains in 1.0, we incorporated 8,298 newly released or updated chains, expanding the dataset to 15,764 chains. Pairwise structural similarity (RMscore) was calculated using RMalign, and all chains were re-clustered to yield 1,042 clusters. Compared with 1.0, 2.0 not only increases sampling density but also identifies 342 new clusters. Notably, Cluster-891 and Cluster-899 correspond to the ROOL RNA and GOLLD RNA families, marking the first inclusion of these large structured bacterial ncRNAs into a global classification framework. Mapping the clusters to Rfam annotated 6,388 chains (40.5%) across 154 families. A single cluster can correspond to one or several Rfam families, indicating that global structure classification and sequence family classification are complementary. RR3DD 2.0 provides an updated web interface.

## Introduction

RNA performs diverse functions such as catalysis, regulation and molecular recognition by folding into specific 3D structures[1]. Studies have shown that the 3D structures of RNA are often more conserved than their sequences, therefore, sequence-based classification alone cannot fully capture the relationship between RNA structure and function[2,3]. Systematic classification of RNA 3D structures is fundamental to understanding the relationship between RNA structure and function[4,5].

Existing RNA classification databases have established relatively mature systems along three levels: sequence, secondary structure, and local 3D structure. At the sequence level, Rfam defines RNA families through multiple sequence alignments and covariance models[6,7], RNAcentral integrates non-coding RNA sequence annotations from multiple specialized databases[8,9], and miRBase[10], GtRNAdb[11,12], NONCODE[13], LncBook[14], SILVA[15] and snoDB[16] further provide specialized classification and annotation by ncRNA functional category. Database resources are continuously updated on platforms such as the NAR Database Issue[17], and comparative genomics has also been used to identify bacterial tRNAs and non-coding RNAs[18]. At the secondary structure level, CRW[19] and RNA STRAND[20] provide comparative sequence and secondary structure information, PseudoBase[21,22] has established a dedicated resource for pseudoknots, the LSU rRNA[23], SSU rRNA[24] and 5S rRNA[25] databases specifically compile rRNA secondary structures, and NNDB[26] provides nearest-neighbor thermodynamic parameters for predicting the stability of nucleic acid secondary structures. At the local 3D structure level, SCOR[4] classifies RNA 3D motifs using a directed acyclic graph, the RNA 3D Motif Atlas[27,28] automatically extracts and clusters hairpin loops and internal loops, RNA FRABASE[29,30] supports 3D fragment retrieval, RNAJunction[31] compiles junctions and kissing loops, RNA CoSSMos[32,33] characterizes secondary structure motifs within 3D structures, and URS[34] compiles RNA structures and their motifs. RNA loop modeling based on sequence homology and geometric constraints[35] and RNA tertiary structure classification methods[5] have also advanced comparative analyses of local and tertiary structures. In addition to classification databases, resources such as RNABase[36], MODOMICS[37], NDB[38], InterRNA[39], RNAInter[40], RNALigands[41], RNApedia[42] and FURNA[43] support RNA structure research from aspects including structural annotation, modifications, base interactions, interactomes and functional annotation, while RNAapt3D[44] provides a resource for 3D modeling of aptamers.

However, none of the above resources has established a classification system at the level of the global 3D fold of entire RNA chains. Classification based on sequences and secondary structures cannot directly reflect 3D folding similarity, and local motif classification cannot address the problem of classifying whole chains at the global structural level. Meanwhile, resources previously involved in RNA structure classification, such as SCOR[4] and DARTS[5], are no longer updated, leaving a gap at the level of global structure classification. By contrast, the protein field has developed a relatively mature global structural classification system, in which fold growth has slowed and structure prediction has made a breakthrough owing to AlphaFold [45,46,58]. Whereas RNA global 3D fold classification still lacks a continuously maintained system, and both RNA structure prediction and experimental structure determination remain in a phase of rapid development [47,48]. This contrast further highlights the necessity of establishing a continuously updatable resource for RNA global 3D structural classification.

To fill this gap, our group established RR3DD in 2021, the first classification database based on global 3D structural similarity of RNA[57]. RR3DD 1.0 collected 13,601 RNA chains from PDB and mmCIF format structures, but its core analysis focused on 7,466 PDB-format chains, which were clustered into 675 RNA folds. The database demonstrated the complementary relationship between global structure classification and sequence family classification: some RNAs that Rfam did not assign to the same family were grouped into the same structural fold in RR3DD, and a single Rfam family could correspond to multiple folds. However, the data in 1.0 were limited to July 2021, after which RNA structure determination entered a period of rapid growth. The application of cryo-EM not only markedly increased the number of structural entries but also enabled the determination of large RNA-only assemblies that previously lacked experimental structures, so that the 1.0 dataset can no longer represent the RNA structure space currently available in the PDB[57]. This growth changed the input to global structure classification in two respects. First, in terms of data volume, the number of RNA structural chains added since 2021 far exceeds the range covered by 1.0. Second, in terms of structural types, the sequences and secondary structure features of ROOL RNA and GOLLD RNA had been reported previously[46,47,48], but both families lacked experimentally determined 3D conformations. In 2025, their 3D structures were determined by cryo-EM[49,50,51,52], providing direct structural evidence for examining whether the 1.0 classification framework covers the fold types of these large assemblies.

Against this background, we established RR3DD 2.0. Starting from the 7,466 PDB-format chains in 1.0[57] and incorporating 8,298 newly extracted chains, we expanded the dataset to 15,764 chains and re-clustered all chains, yielding 1,042 clusters. Compared with 1.0, 2.0 not only increases the sampling density of the structure space but also identifies new clusters absent from 1.0. Among them, Cluster-891 and Cluster-899 are composed of the ROOL RNA and GOLLD RNA families, respectively, giving these two classes of large bacterial ncRNAs independent classification assignments at the level of global RNA 3D structure.

## Materials and Methods

RR3DD 2.0 was systematically expanded from version 1.0[57]. We downloaded 3,626 RNA containing structure files from the Protein Data Bank (released as of 11 November 2025), corresponding to structures newly released or updated after RR3DD 1.0. After removing RNA chains shorter than 5 nucleotides, we extracted 8,298 novel RNA chains not included in RR3DD 1.0. Combining these 8,298 chains with the 7,466 chains in RR3DD 1.0 yielded a final dataset of 15,764 RNA chains.

RNA structural similarity was calculated using RMalign[57], with RMscore as the similarity measure, a higher score indicates greater structural similarity. For the 8,298 newly added chains, we calculated 34,424,253 pairwise scores among themselves and 61,952,868 scores between them and the original 7,466 chains, totaling 96,377,121 new RNA pairs similarity scores. Similarity scores among the original chains were taken from RR3DD 1.0, and a 15,764 × 15,764 similarity matrix was thus constructed.

Clustering was performed using average linkage hierarchical clustering, with the distance defined as d=1−RMscore and the clustering cutoff set at RMscore=0.45. This threshold follows the clustering criterion of RR3DD 1.0 and is based on the RMscore cutoff at which RMalign achieved the highest MCC(MCC = 0.73) in RNA functional classification in previous work[45,53]. RR3DD 2.0 re-clustered all 15,764 chains, the Fold numbering of 1.0 was discontinued, and a total of 1,042 clusters were obtained. Unlike RR3DD 1.0, the Cluster numbering in 2.0 was uniformly assigned after re-clustering all 15,764 chains. Therefore, the Clusters in 2.0 do not correspond one-to-one to the Folds in 1.0 (see Table S2).

## Results

### Cluster statistics and annual variation

Figure 1 shows the member count distribution of the 53 clusters with more than 30 members. Among all 1,042 clusters, the largest, Cluster-1, contains 2,967 chains, and 335 clusters contain only a single chain (Table S1). 84.07% of the clusters contain no more than 5 chains, indicating that most structural types remain relatively rare in the current database. The top 20 largest clusters account for 64.3% of all chains. The chain with the smallest average distance to other members within a cluster was taken as the centroid structure of that cluster. After clustering, each cluster was assigned a unique identifier (Cluster-1 to Cluster-1042), which was used in subsequent analyses.

**Figure 1.**
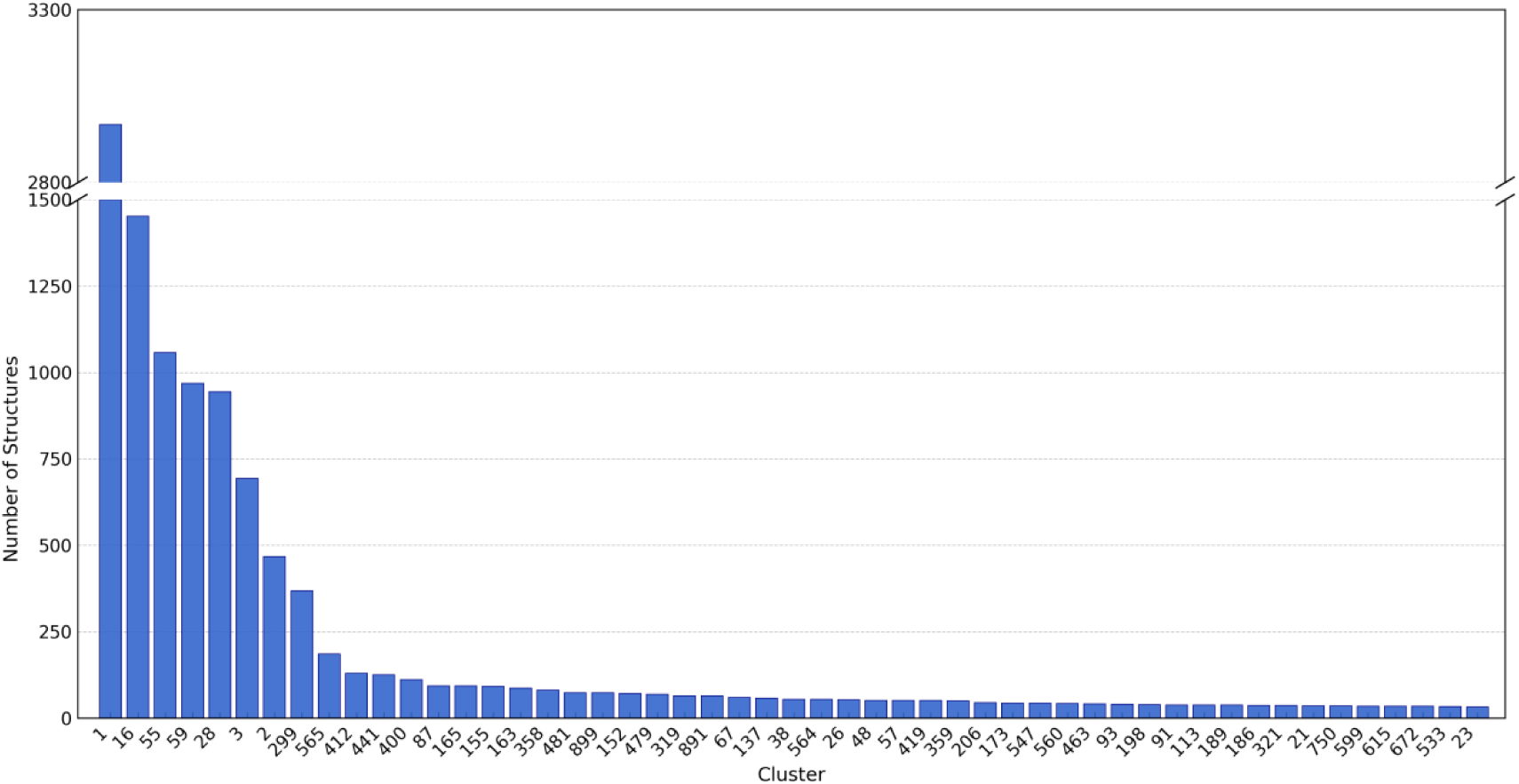
Member distribution of the 53 clusters with more than 30 members in RR3DD 2.0. The x-axis shows Cluster IDs (sorted in descending order of member count), the y-axis shows the number of RNA chains in each cluster. The y-axis is broken between 1,500 and 2,800 to accommodate the largest cluster, Cluster-1 (2,967 chains).

In addition, we counted the number of newly added clusters per year and the cumulative total in RR3DD 2.0 (Figure 2, Table S3). The results show that the expansion of RNA structure space was relatively gradual before 2021. Since 2021, the number of newly added clusters has increased markedly, with 134 new clusters in 2024 and 92 in 2025 (as of 11 November), reflecting the data explosion in recent years brought about by RNA 3D structure determination, particularly through the application of cryo-EM. By the end of 2025, the total number of clusters covered by RR3DD 2.0 had reached 1,042.

**Figure 2.**
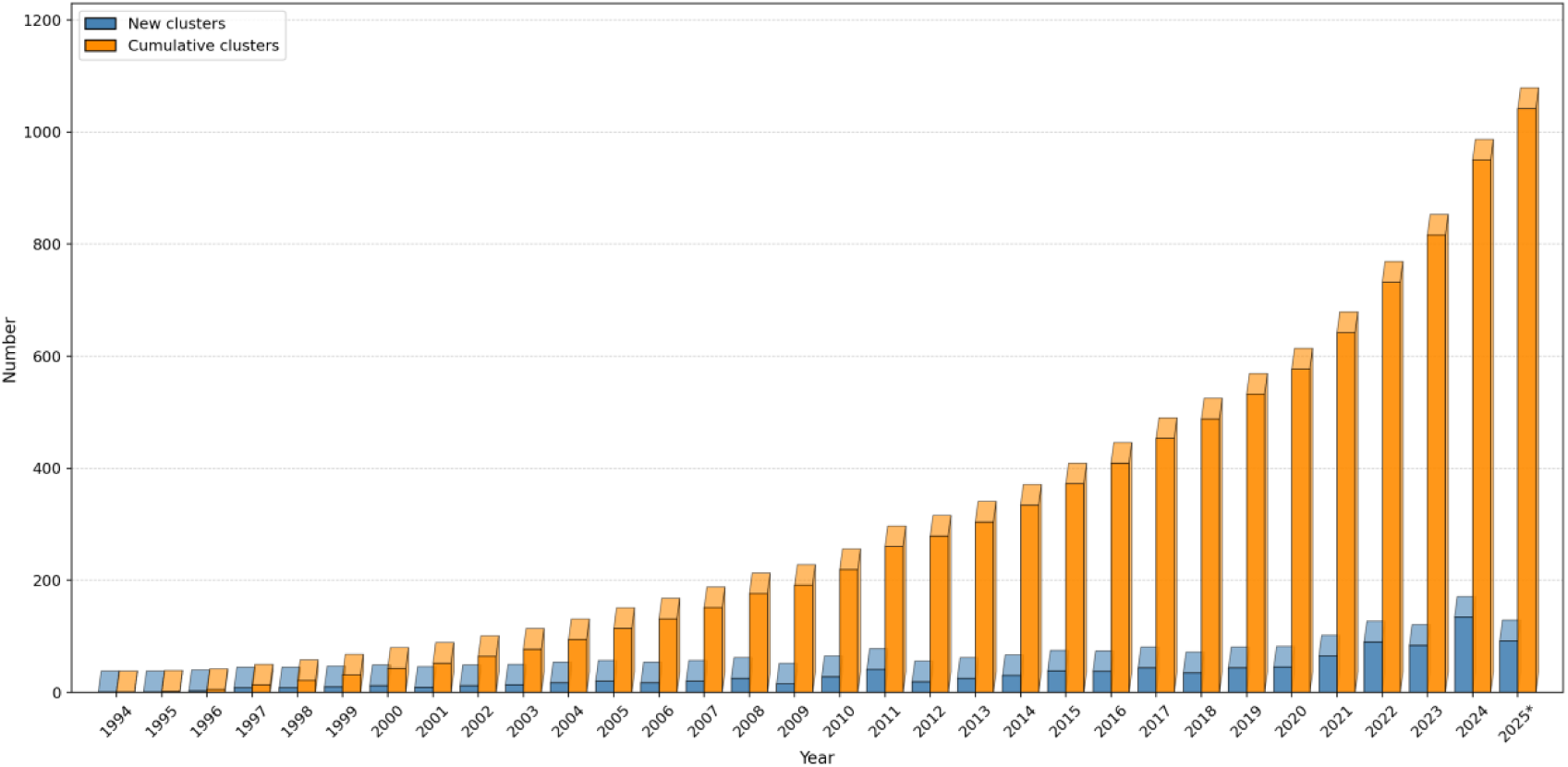
Annual variation in the number of clusters in RR3DD 2.0. The x-axis shows the year, the y-axis shows the number of newly added clusters per year (blue) and the cumulative total number of clusters up to that year (orange). Data for 2025 are as of 11 November 2025.

### Global characteristics of the clustering results Three modes of cluster evolution

To assess how well RR3DD 2.0 covers the RNA structure space, we used principal component analysis (PCA) to visualize the clustering results in reduced dimensions and compared them with those of RR3DD 1.0 (Figure 3, Figure S1). Comparison of the reduced-dimension visualizations of 1.0 and 2.0 showed that the structure space covered by 2.0 is markedly expanded, and that members of most clusters with larger member counts are concentrated within a compact region of the PCA space. Compared with the sparse distribution in 1.0, the RNA structure space in 2.0 appears more compact and continuous. In the distal region of the positive PC1 axis, which was sparsely sampled in 1.0, 2.0 identified several novel clusters, including Cluster-891 and Cluster-899. Figure S1 highlights the distribution of RNA chains newly added in 2.0 relative to 1.0 in the reduced-dimension plot.

**Figure 3.**
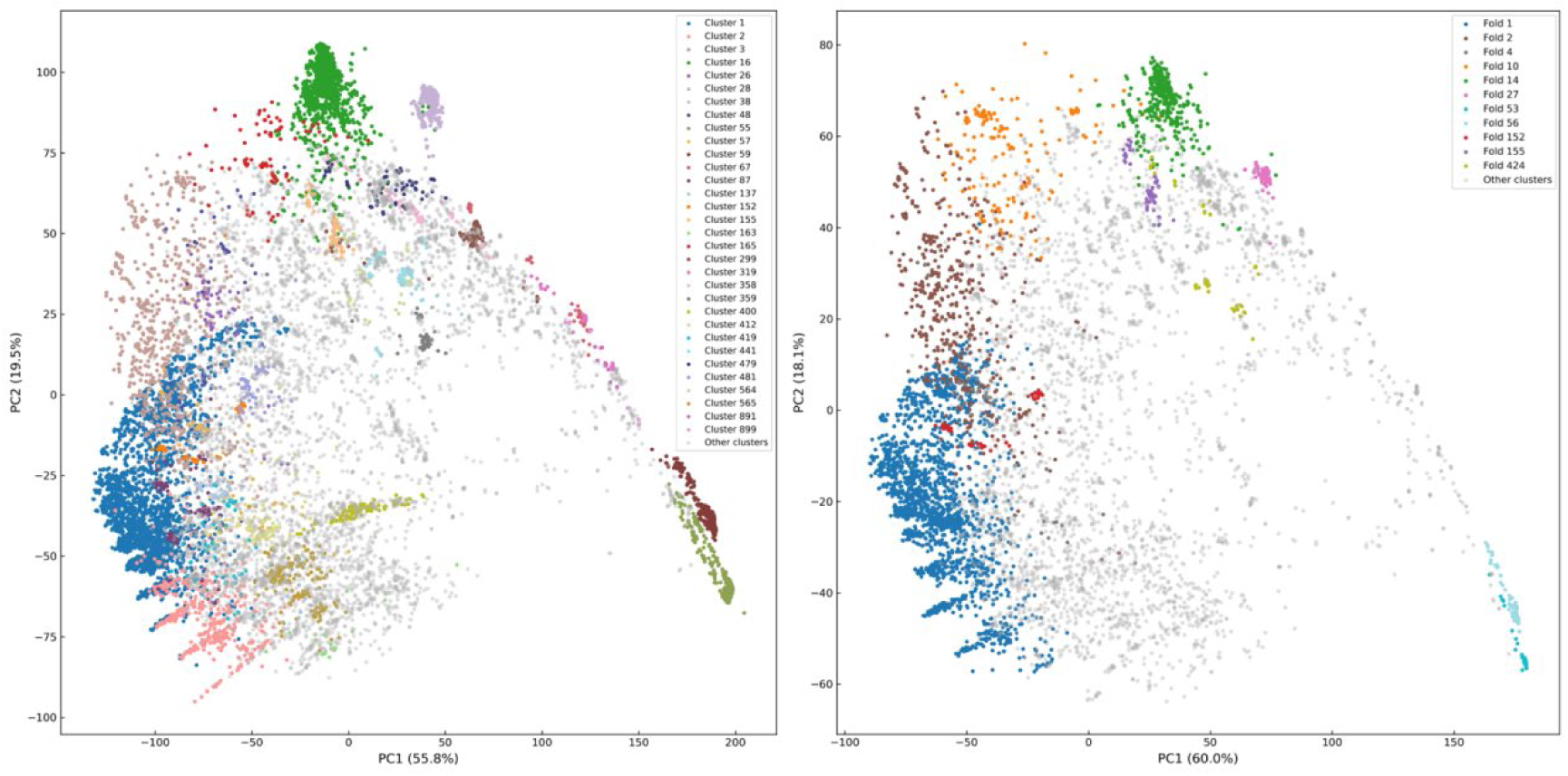
Comparison of PCA visualizations between 2.0 and 1.0. Left, RR3DD 2.0. Right, RR3DD 1.0. Clusters with more than 50 members are colored, with different colors representing different Clusters (2.0) or Folds (1.0), clusters with no more than 50 members are shown in gray.

Cluster are classified evolution into three modes. The first comprises novel clusters, representing structural types that are entirely absent from 1.0 and contributed solely by the newly added 2.0 data. Among the 1,042 clusters, 342 are novel. The two largest novel clusters, Cluster-891 and Cluster-899, correspond to the ROOL RNA and GOLLD RNA families, respectively, and are analyzed in detail in Section 4.

The second comprises conserved clusters, which already have corresponding Folds in 1.0 and acquire few or no new chains in 2.0, so that their clustering results remain stable. For example, Cluster-152 contains 71 chains, all from Fold-152 of 1.0, Cluster-155 contains 92 chains, of which 75 (81.5%) are from Fold-155 of 1.0 and 17 are newly added in 2.0. Taking Cluster-155 as an example, its centroid structure is 6UC8_B (Guanine riboswitch, 67 nt, RF00167). Within the cluster, 62 chains (67.4%) are Purine riboswitches (RF00167), 16 (17.4%) are 2dG-I riboswitches (RF01510), and 14 (15.2%) are not annotated by Rfam. The unannotated members include several Squash RNA aptamers, whose sequence identity to the centroid structure is as low as 0.35 while their structural similarity ranges from 0.55 to 0.80. This phenomenon indicates that Cluster-155 represents a structurally similar RNA fold type rather than a single functional family, showing that the clusters in 2.0 can capture structural similarity across sequence families.

The third comprises expanded clusters, representing structural types whose sampling density increased markedly after the 2.0 data expansion. For example, Cluster-16 expanded from Fold-14 of 1.0 (419 chains) to 1,452 chains in 2.0, with 1,027 newly added, Cluster-28 expanded from Fold-27 (170 chains) to 944 chains, with 774 newly added. Taking Cluster-299 as an example, the cluster contains 368 chains, of which only 12 are from 1.0 (from Fold-288 and Fold-544, respectively) and 356 are newly added in 2.0, the number of chains in 2.0 is 30.7 times that in 1.0. The centroid structure of this cluster is 9B0N_C (5.8S rRNA, 156 nt, RF00002). Of the 368 chains in the cluster, 353 (95.9%) are 5.8S rRNA (RF00002) and the remaining 15 are not annotated by Rfam. For the unannotated members, structural similarity to the centroid structure ranges from 0.44 to 0.86, sequence identity from 0.38 to 1.00, and RMSD from 0.60 to 1.69 Å.

Structural superpositions of the unannotated members of the four representative clusters (Cluster-16, Cluster-28, Cluster-299, and Cluster-155) with their centroid structures are shown in Figure S2, and the corresponding structural similarity, sequence identity, and RMSD values are listed in Table S4.

### Global statistics of Rfam functional annotation

To investigate the relationship between RNA structure and function, we mapped the clusters of RR3DD 2.0 to the Rfam database (Rfam 15.1). The mapping procedure was as follows: two files, pdb_full_region.txt.gz and Rfam.pdb.gz, were downloaded from the Rfam website, where Rfam.pdb provides the mapping from PDB chain identifiers to Rfam IDs and pdb_full_region provides the corresponding region annotations. A custom Python script (Can be downloaded from data materials) was used to associate the PDB chain identifier of each RNA chain in RR3DD 2.0 with its Rfam annotation (Table S1).

The mapping results show that 6,388 of the 15,764 chains (40.5%) received Rfam family annotations, covering 154 different Rfam families. Among the 16 clusters with more than 30 members, some are composed of a single Rfam family (e.g., Cluster-28), whereas others involve multiple families (e.g., Cluster-16) (Figure 4). Cluster-28 contains 944 chains, of which 937 received Rfam annotations, all from a single Rfam family (RF00001). Cluster-16 contains 1,452 chains, of which 1,430 received Rfam annotations covering 9 different families (1,384 of which are tRNAs, RF00005), while 22 chains were not annotated by Rfam, this cluster may reflect the clustering of multiple sequence families under structural constraints. Figure 4 shows the Rfam family composition of the top 10 clusters with more than 30 members.

**Figure 4.**
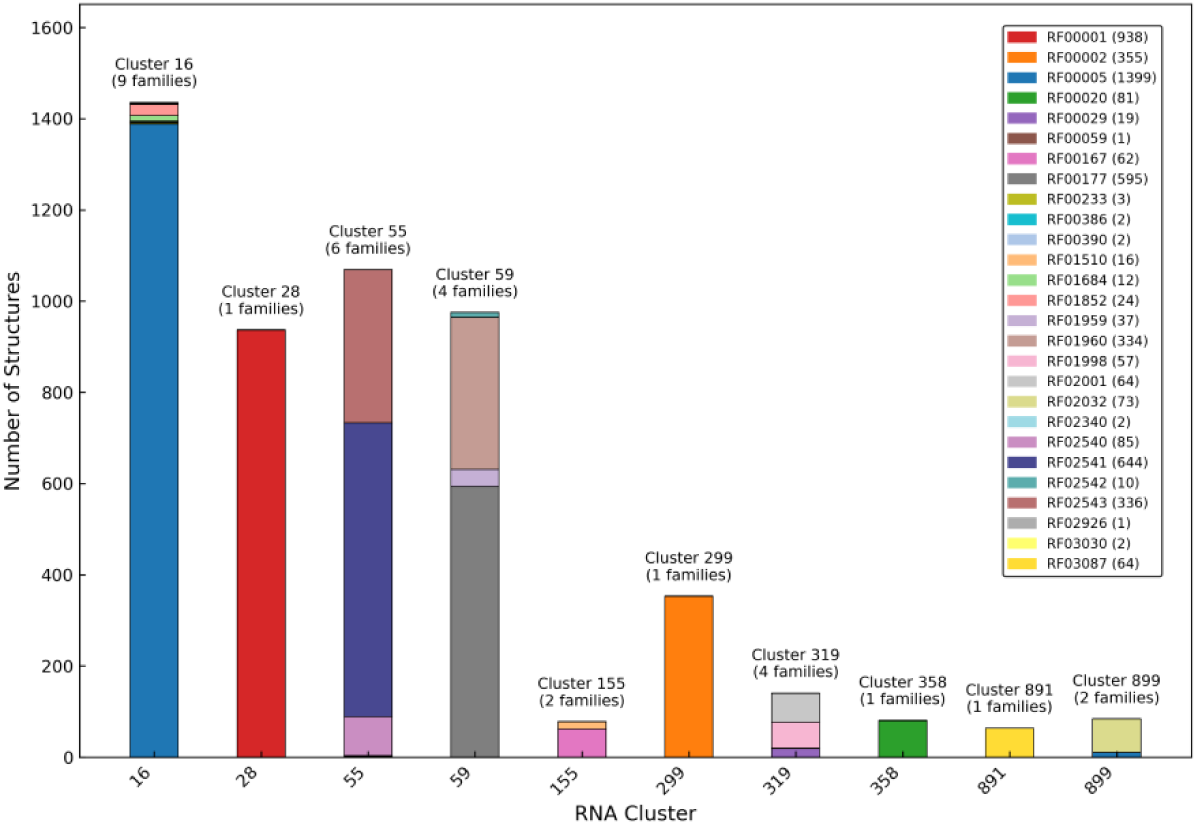
Rfam family composition of the top 10 clusters with more than 30 members in RR3DD 2.0. The x-axis shows Cluster IDs, the y-axis shows the number of RNA chains in each cluster. Different colors represent different Rfam families. A single cluster can map to multiple Rfam families, and the number above each bar indicates the number of Rfam families corresponding to that cluster.

### Sequence-structure relationship analysis

To investigate the relationship between sequence conservation and structural conservation of RNA, we analyzed the distribution of sequence identity and structural similarity (RMscore) for the 96,377,121 newly added RNA pairs (Figure 5). The sequence identity threshold of 0.6 and the RMscore threshold of 0.45 follow the analysis criteria of RR3DD 1.0[57]. When sequence identity is below 0.6, the relationship between sequence and structural conservation weakens, and 0.45 is the transition point between similar and dissimilar RNA 3D structures. The distribution across the four quadrants is shown in Table 1.

**Figure 5.**
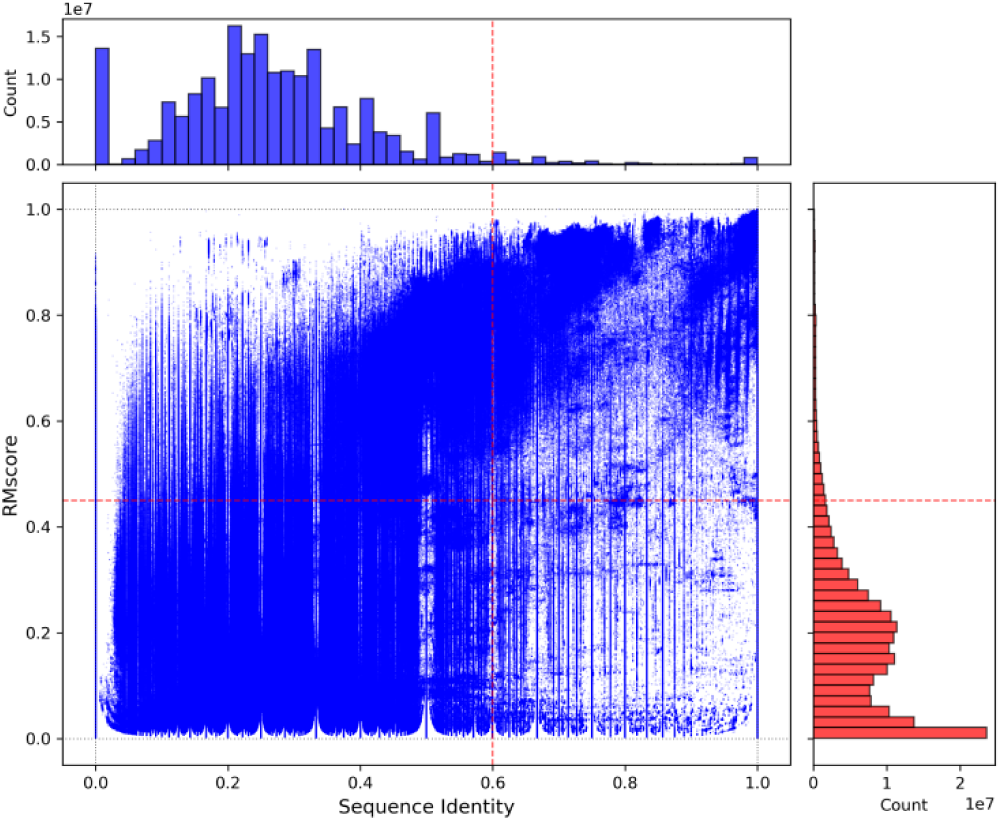
Relationship between RNA sequence identity and structural similarity (RMscore). The x-axis shows sequence identity, the y-axis shows RMscore. Dashed lines indicate the sequence identity threshold of 0.6 and the structural similarity threshold of 0.45, dividing the data into four quadrants. The marginal histograms at the top and on the right show the frequency distributions of sequence identity and RMscore, respectively.

**Table 1.** Quadrant distribution of sequence identity and structural similarity (RMscore) for the 96,377,121 newly added RNA pairs.

| Quadrant | Definition | Number | Proportion |
|---|---|---|---|
| Q1 | sequence identity > 0.6, RMscore > 0.45 | 960,493 | 1.00% |
| Q2 | sequence identity ≤ 0.6, RMscore > 0.45 | 5,214,178 | 5.41% |
| Q3 | sequence identity ≤ 0.6, RMscore ≤ 0.45 | 88,857,359 | 92.20% |
| Q4 | sequence identity > 0.6, RMscore ≤ 0.45 | 1,345,091 | 1.39% |

The vast majority of RNA pairs (92.20%) have both low sequence identity and low structural similarity, consistent with the 93.42% observed in 1.0. The Q2 quadrant (low sequence identity, high structural similarity) contains 5.41% of the RNA pairs, close to the 5.44% in 1.0, the structural similarity of these pairs cannot be explained by sequence similarity alone, indicating that structural alignment can identify similar structures missed by sequence alignment. The Q1 quadrant (high sequence identity, high structural similarity) accounts for 1.00%, close to the 1.02% in 1.0. Notably, the Q4 quadrant (high sequence identity, low structural similarity) accounts for 1.39% in 2.0, markedly higher than the 0.12% in 1.0, indicating that RNA pairs with similar sequences but dissimilar structures are not rare in the expanded data, this phenomenon may be related to conformational polymorphism or different folding states.

### Case studies of novel clusters: ROOL RNA and GOLLD RNA Cluster-891 (ROOL RNA)

Cluster-891 is a global 3D structure cluster newly identified in RR3DD 2.0. It is located in the distal region of the positive PC1 axis in the reduced-dimension space and contains 64 RNA chains with sequence lengths ranging from 520 to 650 nucleotides. All 64 chains are annotated as the Rfam family RF03087 (ROOL RNA), indicating high consistency of this cluster at the sequence family level.

In 2017, Weinberg et al. predicted the existence of this non-coding RNA through comparative genomic analysis of rumen metagenomic data and described a complex secondary structure containing 6 multistem junctions and 3 pseudoknots, with a predicted average length of 581 nt[54]. In the same year, Cousin et al. independently identified a highly expressed lncRNA of 520 nt by RNA-seq in Lactobacillus salivarius UCC118, whose abundance in stationary phase can exceed that of 16S rRNA[55], this lncRNA was later identified as ROOL RNA[56]. Neither study obtained a 3D structure of ROOL RNA. In 2025, cryo-EM structures revealed the 3D conformation of ROOL RNA for the first time, showing that it can assemble into hexameric[54] or octameric[57] nanocages.

Using 9LE5_A as the reference structure, pairwise structural alignment revealed substantial heterogeneity within this cluster. RMSD values range from 0.27 to 5.04 Å, and all RMscore values are above 0.5. The 64 RNA chains in Cluster-891 cover multiple oligomeric states of ROOL RNA (Table S5). Among them, 4 chains exist as monomers (9ISV_A, 9LE5_A, 9M79_A, 9MMG_c), while the remaining 60 chains are subunits of tetramers (9J3R), hexamers (9LE3, 9LE6, 9J6Y, 9M78), and octamers (9J3T, 9MM6, 9MME, 9MDS). The monomeric units of these different oligomeric states are highly similar in global shape, and Cluster-891 simultaneously captures the monomeric subunits of these different oligomeric states, indicating that the global fold of the ROOL RNA monomer is conserved across different assembly environments, although the local conformation is adjusted with the oligomeric state.

The subunits corresponding to different multimeric states differ in RMSD (0.27– 5.04 Å); hexameric subunits are more similar to each other (RMSD < 1.74 Å), whereas octameric subunits differ more from the reference structure 9LE5_A (monomer) (RMSD 4.1–5.0 Å). This observation may reflect the conformational adaptability of ROOL RNA under different assembly states and may be related to its ability to form different multimeric forms.

ROOL RNA has unusually high structural stability, degrading by only about 10% within 30 min after transcription inhibition, a stability comparable to that of 16S rRNA[55], which may be related to its highly conserved global fold. This RNA remains expressed during colonization of the mouse intestine by L. salivarius[55], suggesting that it may function in host–microbe interactions.

### Cluster-899 (GOLLD RNA)

Cluster-899 is likewise located in the distal region of the positive PC1 axis. It contains 73 RNA chains, all annotated as the Rfam family RF02032 (GOLLD RNA). This cluster is also absent from RR3DD 1.0.

GOLLD RNA was first discovered in 2009 through bacterial metagenomic analysis and is the third largest known structured bacterial non-coding RNA, after 23S and 16S rRNA[56]. In 2025, Zhang et al. resolved the high-resolution structure of GOLLD RNA by cryo-EM for the first time, revealing that full-length Sag-GOLLD RNA forms a dodecamer and Env-38 GOLLD forms a tetradecameric nanocage[54]. Cluster-899 uses 9LEE_J (a Sag-GOLLD RNA monomer) as the centroid structure. Pairwise structural alignment using 9LEE_J as the reference shows that RMscore values within the cluster range from 0.48 to 1.00, sequence identity from 0.49 to 1.00, and RMSD from 0 to 2.66 Å (see Table S5).

Notably, 10 chains in Cluster-899 carry both GOLLD and tRNA annotations (RF00005), corresponding to the Sag-GOLLD-pretRNAser fusion RNA, in which the tRNA portion is linked to GOLLD RNA through a 10-nucleotide linker[54]. According to PDB annotations, 9MEE and 9LHL form tetradecamers, 9LMF, 9LEE, and 9LEM form decamers, and 9L0R forms dodecamers, all 73 chains in Cluster-899 are monomeric subunits of these multimers.

Cluster-891 (ROOL RNA) and Cluster-899 (GOLLD RNA) are two novel clusters in RR3DD 2.0 contributed entirely by newly added data. Both are located in the distal region of the positive PC1 axis in the reduced-dimension plot and represent shape space not previously covered by other RNA structure classification database. The identification of these two clusters incorporates large bacterial ncRNA families (ROOL and GOLLD) into the global 3D structure classification framework for the first time, bridging sequence-based predictions of ROOL and GOLLD and their 3D structural validation.

## Discussion

RR3DD 2.0, established in this work by incorporating recently added RNA 3D structures and re-clustering, not only expands the coverage of the structure space but also captures novel fold types that were previously unclassified. The emergence of Cluster-891 (ROOL RNA) and Cluster-899 (GOLLD RNA) indicates that the cryo-EM-driven determination of large RNA-only assemblies is continuously extending the boundaries of the global RNA structure space. By incorporating these newly identified structures into the global classification framework, they can be observed in a comparable structural space, which is difficult to directly provide for classification based on sequences and secondary structures. The convergence of multiple families (mainly tRNAs) and unannotated members within the same structural cluster in Cluster-16 further corroborates the complementarity between structural classification and sequence family classification: a single structural fold can accommodate members with substantial sequence differences. However, the methodological framework of 2.0 has limitations: clustering operates on individual RNA chains, and the higher-order architectural information of multimeric assemblies is not directly encoded in the clustering features. How to incorporate assembly-level structural differences into the classification framework without compromising the consistency of the existing classification is a question that future versions of RR3DD will need to address.

## Acknowledgements

This work has been supported by the National Natural Science Foundation of China ([32271267] and [32571416]), and HPC Platform of Huazhong University of Science and Technology.

## Author contributions

Conceptualization, DYL and SYL; Investigation DYL; RR3DD software development, DYL and SYL; Data Curation, DYL; Writing - Original Draft, DYL; Writing - Review & Editing, DYL, XDL, CGL, YCZ and SYL; Funding Acquisition, SYL; Supervision, SYL.

## Data availability

pdb_full_region.txt.gz and Rfam.pdb.gz can be downloaded at http://ftp.ebi.ac.uk/pub/databases/Rfam/CURRENT/database_files/pdb_full_region.txt.gz and http://ftp.ebi.ac.uk/pub/databases/Rfam/CURRENT/Rfam.pdb.gz, respectively. The RR3DD2.0 webpage can be found at http://www.rnabinding.com/RR3DD2.0/.

